# The Application of The Skin Virome for Human Identification

**DOI:** 10.1101/2021.09.10.459834

**Authors:** Ema H. Graham, Jennifer L. Clarke, Samodha C. Fernando, Joshua R. Herr, Michael S. Adamowicz

## Abstract

The use of skin virome for human identification purposes offers a unique approach to instances where a viable and statistically relevant human DNA profile is unavailable. The human skin virome may act as an alternative DNA profile and/or an additional form of probative genetic material. To date, no study has attempted to investigate the human virome over a time series across various physical locations of the body to identify its potential as a tool for human identification. For this study, we set out to evaluate the stability, diversity, and individualization of the human skin virome. An additional goal was to identify viral signatures that can be used in conjunction with traditional forensic STR loci. In order to accomplish this, human virome metagenomes were collected and sequenced from 42 individuals at three anatomical locations (left hand, right hand, and scalp) across multiple collections periods over a 6-month window of time. Assembly dependent and independent bioinformatic approaches were employed, along with a database-based assessment, which resulted in three sets of stable putative viral markers. In total, with the three sets combined, 59 viral species and uncharacterized viral genome assemblies were identified as being significantly stable (*P*=5.3×10^-15^). Viral diversity, based on presence or absence, is significantly different across subjects (*P*<0.001). Here we demonstrate that not only is the human virome applicable to be used for human identification, but we have identified many viral signatures that can be used for forensic applications, thus providing a foundation to the novel field of forensic virology.

**Highlights:**

- Here we provide the largest human skin virome study, to date. Our study revealed novel diversity findings of high abundance for certain viral taxa, for example, the Cress-like DNA phages, that have not previously been characterized in human skin viral ecology studies.
- There were 59 putative human skin viral biomarkers suitable for human identification from the core stable human skin virome of 42 subjects.
- The putative markers we identified were significantly stable over a 6-month period of time within individuals and across three autosomal locations of left hand, right hand, and scalp.
- Diversity of profiles, based on the presence and absence of our putative marker data set, were significantly different across test subjects.

## 1. Introduction

The use of microbial signatures for human identification constitutes a new frontier in forensic applications. The wide variety of data types, sample locations, environmental factors, analysis methods, and the diversity of microorganisms make this emerging area of forensic biology increasingly more relevant [1]. The potential of utilizing the human microbiome in forensic science has previously been described by Hampton-Marcell *et al* [2] who performed experiments to identify associations between the human skin bacterial composition and the physical environment. That study concluded that using 16S rRNA sequencing of bacteria provided weak predictions of human identity compared to more traditional methods of forensic human DNA analysis. However, they also recognized that there could be better methods of analysis and that the field is currently in its infancy [2]. Although the application of microbiome analysis in relation to casework is relatively new, the use of bacteria in forensically relevant cases dates back to the late 1800s [3]. In one modern example, researchers have examined commercial honey products for the presence of *Clostridium botulinum* as a vector for cases of induced poisoning from botulism [4]. More recently, *Neisseria gonorrhoeae* infections in children have been considered as evidence in sexual assault cases [5]. As such, the application of microbes in forensic applications is not a new concept.

With the advancement of sequencing technologies and decreased sequencing costs, the study of meta-communities of microorganisms for forensic applications began to be addressed in earnest with a series of studies identifying and defining the discipline [6–10]. Since then, the discipline of forensic microbiology has progressively expanded. For example, Tims *et al* [11] investigated the possibility of isolating microbial DNA from human fingerprints. They concluded that the bacterial microflora on human fingertips was too dynamic to develop reliable markers and thus not suitable for human identification. Expanding on that study, recent work using massively parallel sequencing and metagenomic analysis have demonstrated that bacterial DNA can be collected from an individual fingerprint in quantities sufficient to create a profile [12]. Those researchers concluded that individual profiles of the human hand microbiome were highly variable [12]. Application of microbiome differences have also been used to explore insoles of footwear and the soles of feet utilizing termination restriction fragment length polymorphisms (T-RFLPs) [13]. This work also indicated that bacterial profiles could be identified from individual people and that the right and left plantar microbiome are significantly similar to each other, while those between people were not [13]. Microbial forensic research has also been applied to the identification of body fluids. For example, Giampaoli *et al* [14] used mixed microbial populations to positively identify vaginal fluid, even when vaginal fluid was mixed with other body fluids. Research that distinguished vaginal from saliva, feces, or skin samples through DNA sequencing and microarray analysis concluded that a combination of microarray and probabilistic analysis would be best suited for determining body site location [15].

Individualization of pubic hair samples using sequence data from bacterial communities was attempted by Williams and Gibson [16] who found that a large variation in microbiome community was observed across gender and individuals, therefore suggesting that it may be possible to use microbiome data for human identification in criminal investigations for crimes such as rape and sexual assault. The same researchers expanded on their work, showing that microbiome samples from the pubic mound area can differentiate individuals and that couples exhibited increased similar microbiomes than those of random people [17]. The differences between the penile and vaginal microbiomes have also been examined, resulting in data demonstrating that the sexes also show detectable microbial signatures that may be used to differentiate people and could be highly relevant in cases of sexual assault where no ejaculation has occurred [18]. The transfer of skin-associated microbiomes to objects has been examined by several groups [19–22] using a variety of processing and analytical methodologies. A common theme that has emerged suggests that results must be interpreted with care due to factors such as background organisms and contamination. Despite these cautionary factors, there is positive evidence suggesting that microbiome forensic approaches have merit.

While there has been ample work over the last two decades using bacteria as targets for forensic applications, there have been very few studies addressing the application of viruses and the human virome for forensic purposes and human identification. A potential method for tracing unidentified human cadavers using the JC virus was published by Ikegaya *et al* [23] where the authors suggested that the presence of this virus in humans may aid in determining the geographical location of origin of the decedent. Wilson *et al* [24] laid out specific considerations and presented seven discrete steps for analysis of viral samples in a forensic case. The authors also described a broad range of topics relevant to working with viral biomarkers, including considering viruses as weapons and the intentional/unintentional transmission of virus-mediated diseases in criminal cases. While not developed with a forensic purpose, some markers, such as *Propionibacterium* phage, may be used as potential discriminating factors, and are also part of hidSkinPlex, a panel of markers which were developed for the assessment of skin microbiome diversity from a human health perspective [21].

To date, no study has attempted to investigate the human virome over a time series across various physical locations of the body to identify its potential as a tool for human identification. One reason for the limited number of virome studies has been the lack of bioinformatic and molecular tools for human virome investigation. Despite this, the development of massive parallel sequencing methods and the resulting decrease in sequencing costs have led to studies investigating the viromes of humans, primarily focusing on describing viral diversity in the human gut [25–30]. These studies have demonstrated that human gut viromes tend to be “highly individual and temporally stable” [29], two key features that are essential for a good biomarker for forensic science.

In order to address the human skin virome in a forensic context, we investigated the temporal human skin virome stability over a 6-month period across three body locations left hand, right hand, and the scalp) in 42 individuals using five longitudinal samples from each participant. Our hypothesis is that, one, the human skin virome consists of both stable and variable viral taxa and, two, identifying unique but stable viral sub-populations within the human virome may provide opportunity to utilize virome diversity for forensic human identification.

## 2. Materials and Methods

### 2.1 Sample Collection

Samples from the skin virome of the participants were collected using a tandem dry and wet swab technique using nylon flocked swabs (4NG Floq Swabs, Copan, Brescia, Italy), as previously described [31, 32]. The wet swab was moistened with sterile 1x phosphate buffered saline (PBS) and acted as the leading swab and was immediately followed by the dry swab to collect loosened sample material. Post swabbing, all collected swabs were stored in pre-sterilized Eppendorf safe lock 2ml tubes (Eppendorf, Hamburg, Germany). Virome samples were collected from 42 adult individuals with ages ranging from 19 to 70+ years, who were not on antibiotic drug regimens during the duration of the sampling. Originally 43 participants were a part of this study, however, one subject (P33) was unable to complete the study due to the required use of antibiotics during the duration of sampling time points and was therefore not used for the analysis of this study. Samples were collected across a longitudinal 6-month period to represent the initial sampling (day 0), and 2-weeks, 1-month, 3-months, and 6-months intervals from the initial sampling date. At each collection, virome swabs were collected from three skin locations - left hand, right hand, and scalp. At each sampling date the participants filled out a questionnaire to gather information on travel, skin care, lifestyle, and other information that could help identify factors affecting the microbiome. A representative questionnaire used in the study is included as Supplementary Fig. 1. During each sampling, negative control swabs were collected to evaluate any contamination that may have resulted from laboratory factors, including the batch of PBS used and DNA extraction date, as well as subsequent sequencing run effects. Forensic samples would most likely not consist of highly degradable RNA, so this initial study focused on DNA viruses, and as such, the collected swabs were stored at -20℃ until used for viral enrichment and sequencing.

### 2.2 Viral Enrichment and Purification

Swabs containing skin virome samples were saturated with 200µl of 0.02µm filtered sterile 1x PBS and were placed in a 2ml tube containing a CW Spin Basket (Promega, Madison, WI, USA). Swabs were centrifuged at 16000xg for 10 minutes to elute viral particles from the swab into the PBS solution. The filtrate containing the viral particles was further filtered using a 0.22µm filter to remove cellular and bacterial contaminants. Enriched for viral particles, the resulting filtrate was used for viral DNA extraction using the QiAmp Ultra-Sensitive Virus Kit (Qiagen, Hilden, Germany) according to the manufacturer’s protocol.

The resulting viral DNA was subjected to whole genome amplification (WGA) using multiple displacement amplification (MDA) implemented with the TruePrime WGA Kit (Syngen Biotechnology, Inc, Taipei City, Taiwan) following the standard product protocol. Following WGA, the samples were quantified using the DeNovix dsDNA High Sensitivity Kit using the DeNovix DS-11 Spectrophotometer/Fluorometer (DeNovix, Inc, Wilmington, DE, USA).

### 2.3 Viral Metagenome Library Preparation and Sequencing

One hundred nanograms of the amplified DNA was used for library preparation. The DNA was sheared using sonication to a mean length distribution of 600 bp. The sonication was performed using a Bioruptor (Diagenode, Denville, NJ, USA) with three cycles of 30 seconds on and 90 seconds off as per manufacturer instructions. The resulting sheared DNA was used for library preparation using the NEBNext Ultra II Library preparation kit (New England Biolabs, Ipswich, MA, USA) according to the manufacturer’s protocol. During the kit process, recommended conditions for size selection of adaptor-ligated DNA approximate insert size of 500-700 bp was used. Followed by PCR enrichment of adaptor ligated DNA with five cycles of denaturation and annealing/extension. Final library preps were evaluated using an Agilent 2100 Bioanalyzer (Agilent Technologies, Inc, Santa Clara, CA, USA) with high sensitivity chips to identify sample base pair distribution and sample concentration. Additionally, libraries were quantified using the DeNovix dsDNA High Sensitivity Kit (DeNovix, Inc, Wilmington, DE, USA). The libraries were then sequenced using the 150 bp paired-end sequencing strategy on the Illumina Hiseq 2500 platform (Illumina, Inc, San Diego, CA, USA). All raw sequencing data has been deposited in the NCBI Short Read Archive (SRA) under the accession code PRJNA754140.

### 2.4 Assembly of Metagenomic Sequencing Reads

The resulting sequencing data in Fastq format quality was evaluated using FastQC v.0.11.9 [33] and trimmed and filtered using the adaptive trimming tool Sickle v.1.3.3 [34] to remove low quality reads using a quality filter threshold of Q30 and a length threshold of 75 bp. Reads resulting from the Phi X spike-in were removed from trimmed reads using BBDuk command with standard operational flags from the BBMap suite of tools [35]. Following quality filtering, bacterial contamination was assessed by mapping trimmed reads to the Silva 16S ribosomal database v.138.1 using BBMap flags and parameters described for high precision mapping of contamination detection as suggested in the BBMap Guide [35]. Additionally, all sample reads were mapped to the human genome (hg19) using BBMap with standard operational flags [35], for high precision mapping with low sensitivity in order to lower the risk of false positive mapping. All mapped reads were removed to ensure the viromes are devoid of contamination from bacterial and human host sequences. Metagenome assemblies were performed using MEGAHIT v.1.2.8 [36]. Assemblies were performed using two approaches, 1) assembly within each sample, and 2) a master meta-assembly using all reads. Assembly quality was assessed using QUAST v.5.0.2 [37]. In addition, a meta-assembly using all negative control reads was performed to identify potential contaminants in the dataset that may arise from reagents. The virome assemblies generated were mapped to the negative control contigs greater than 1000 bp using BWA-mem [38] to remove any reads that may have resulted from contamination. Subsequent contigs greater than 1000 bp were then utilized for downstream analysis of viral identification, diversity analysis, and assessment of stability of the virome.

### 2.5 Viral Identification and Taxonomic Classification

Putative viral contigs -- designated as contigs containing annotated viral genes -- were identified using the tool CheckV v.0.7.0 [39]. Contigs first identified as viral via CheckV were additionally subjected to other classification schemes using various viral annotation and classification tools as described below. Viral contigs were further classified using both nucleotide-based classification tools, such as Kraken2 v.2.0.8-beta [40], Demovir [41], and Blastn (with a >10% query coverage cut-off) [42], and also using a classification tool based on protein coding sequences, Kaiju v.1.7 [43]. The least common ancestor of the consensus hits was used for classification results having similar e-values or percent confidences but different taxonomic results to reduce misclassification of viral contigs. With all of the classification tools, the resulting classification we used moving forward was based on those hits having the lowest e-value and/or highest percent confidence.

### 2.6 Mapping of Raw Reads to Assemblies and Viral Contigs

The raw sample data, in the form of forward and reverse paired-end reads, were mapped to the metagenome assembly consisting of contigs from the sample-by-sample based assembly. Read mapping was then performed using Bowtie 2 v.2.3.5 [44] and SAMtools v.1.9 [45] to identify the abundance of each viral contig within each of the respective samples. The resulting read abundances were further analyzed using “R” v.3.6.3 [46]. Unique contigs were used to identify viral diversity and temporal changes. Using the phyloseq R package, a phyloseq object table was created in R using the read abundance data and this table was subsequently used for diversity analysis [47]. Annotation of viral contigs was performed using the aforementioned classification tools described above and incorporated in the phyloseq object table. The count table and mapping file information from the participant questionnaire was used as input for phyloseq object generation, allowing for both *denovo* and reference-based analysis of the skin virome data.

### 2.7 Assessment of Virome Stability Over Time

Viral contig stability was assessed on the basis of presence and absence of viral contigs over time across each body site sampled (left hand, right hand, or scalp) within each subject. Contigs present in four out of the five time points from a specific location within an individual were considered to be a stable viral contig and were identified as a potential marker for human identification. Out of the identified viral contigs, those that were not identified on both hands of a given individual were removed. Viral contig stability was assessed at the taxonomic level of family, genus, and species.

An assembly independent method was also employed using the identified stable viral families for further refinement of viral taxonomic identification at the species and genus level and investigation into putative human identification markers. Based on the preliminary taxonomic assessment of the most abundant and stable viruses, the corresponding reference sequences to the viral families *Papillomaviridae*, *Genomoviridae*, *Baculoviridae*, as well as the order *Caudovirales* were downloaded from the NCBI nucleotide database for subsequent analysis. Raw sequencing reads were mapped to the reference sequences identified as belonging to using Bowtie 2 v.2.3.5 [44]. The resulting mapped reads to the database reference sequences were acquired by filtering with SAMtools [45]. The sequencing data and read counts identified at this stage were utilized for further investigation of viral diversity and persistence of selected viral families and for statistical evaluation using R. To statistically assess stability of the identified markers a Jaccard dissimilarity matrix was produced with the option of binary set to true, with the resulting distance of 0 being all classes being the same and 1 being all classes being disjointed. The subsequent Jaccard distances were averaged between a subject’s sample and all other samples (between dissimilarity) and averaged all distances between a subject’s sample and all other samples collected from that same subject (within dissimilarity). We also calculated Bray-Curtis similarity for the samples, but we have focused on Jaccard in this study because it is more discriminating for samples consisting of presence-absence data [48] and the Bray-Curtis calculation did not fundamentally differ from the Jaccard analysis. Pairwise Wilcoxon rank sum tests were performed between the within subject dissimilarity distances and the between subject dissimilarity distances to assess stability of the markers.

### 2.8 Viral Metagenome Diversity

Alpha diversity (ɑ-diversity) was assessed using the Shannon alpha diversity metric [49]. To evaluate if the contig diversity of a subject significantly changed across body sites and time, a one-way ANOVA using repeated measures was used. For Beta diversity (β-diversity), a binary Jaccard dissimilarity matrix with the option of binary set to true was generated based on presence and absence of identified putative viral human identification markers. Beta diversity was visualized using a principal coordinate analysis (PCoA). The subsequent binary Jaccard dissimilarity matrix was used for PERMANOVA analysis using the Adonis test in the Vegan package v.2.5-7 [50] to assess changes of the virome between subjects. These analyses were all conducted using the R statistical package and using the R associated tools Phyloseq and Vegan [46,47,50]. All metadata, list of markers, contig sequences, annotation files, and scripts described in the material and methods are publicly available and archived at: https://github.com/HerrLab/Graham_2021_forensics_human_virome.

## 3. Results

### 3.1 Virome Assembly and Annotation

Skin virome samples were collected from 42 individuals across three locations (left hand, right hand, and scalp) over a six-month period - with samples representing the initial timepoint (day 0), 2-weeks, 1-month, 3-month, and 6-month timepoints taken from the initial sampling date. These physical locations were chosen based on their potential to be of forensic relevance, level of viral diversity, and microbiome temporal stability as shown in our pilot sampling and previously published studies [28, 51].

Samples were pre-processed using 0.2-micron filtration to remove eukaryotic and prokaryotic cells and post-processed with bioinformatic analysis to remove reads with high synteny to the human genome. The resulting quality filtered reads were used to evaluate viral diversity and viral stability with the long-term goal of identifying viral markers for human identification. Out of the read assemblies only contigs larger than 1000 bp were used for subsequent analysis. In total, out of the 952,760 contigs, 62,101 contigs were >1000 bp and thus retained for further analysis. This resulted in the contigs having a N50 value of 1970 and the longest contig length being 54,929 bp.

To ensure that the viral contigs were devoid of bacterial contamination, sample assemblies were evaluated for the presence of 16S rRNA genes. Sample metagenomic assemblies contained on average 0.002% of ribosomal reads per sample and a maximum of 0.16% of ribosomal reads. Previous studies have demonstrated that if viral metagenomes have less than 0.2% 16S rRNA reads, these datasets are enriched for viral sequences and have minimal and likely negligible bacterial contamination [52]. As such, the low occurrence of bacterial 16S rRNA sequences suggests that our viromes were adequately enriched for human-associated viruses with minimal contamination. To further identify contigs of viral origin based on currently available viral sequence information, contigs were further analyzed using CheckV v.0.7.0 [39]. Out of the 62,101 assembled contigs, 1400 were identified as having a known viral gene and assumed to be truly viral in origin based on current databases [39]. Of the 1400, 102 contigs were removed for having 100% synteny to the human or animal (e.g., *Canis lupus*) genomes or considered prophage, resulting in 1298 final contigs. Our viral metagenome dataset is the largest dataset to date from the human skin virome and may contain many novel viruses that cannot be annotated using the current databases. Additionally, as described above, our viral metagenomes are devoid of bacterial contamination as such we believe we have a significant portion of uncharacterized novel human viruses contained in our dataset.

### 3.2 Human Skin Viral Diversity

The 1298 identified contigs were taxonomically classified using current viral databases and viral annotation bioinformatic tools and the distribution of abundance by taxonomy of the contigs across all samples is represented in Fig. 1. A majority of the identified viral contigs could not be fully annotated using currently available viral reference databases and existing bioinformatic tools. Even the contigs that were identified as viral contigs through CheckV, could not be classified to lower taxonomic levels using currently known reference viral genomes suggesting that existing public viral databases are poorly representative of human skin viral diversity leading to taxonomic uncertainty for many contigs in our study. Due to the fact that we sequenced the DNA virome, our contigs show synteny to double and single stranded DNA viruses. However, many of the DNA viruses identified were unclassified due to no representation in the current viral databases. Many such viruses were identified as highly abundant in the core human skin virome, as shown in Fig. 1. Among the double stranded DNA viruses identified, the viral order *Caudovirales* was the most abundant order detected (Supplementary Fig. 2). This is not surprising, as there is a disproportionate amount of *Caudovirales* viral genomes available in viral reference databases. As for the identified single stranded DNA viruses, highlighted in Supplementary Fig. 3, the most abundant taxa were that of small circular DNA viruses which included Papillomaviruse*s* and Cress-like DNA phages. Papillomaviruses and Polyomaviruses are common skin associated opportunistic pathogens and the identification of Papillomavirus genomes was expected [28, 53].

**Fig. 1.**
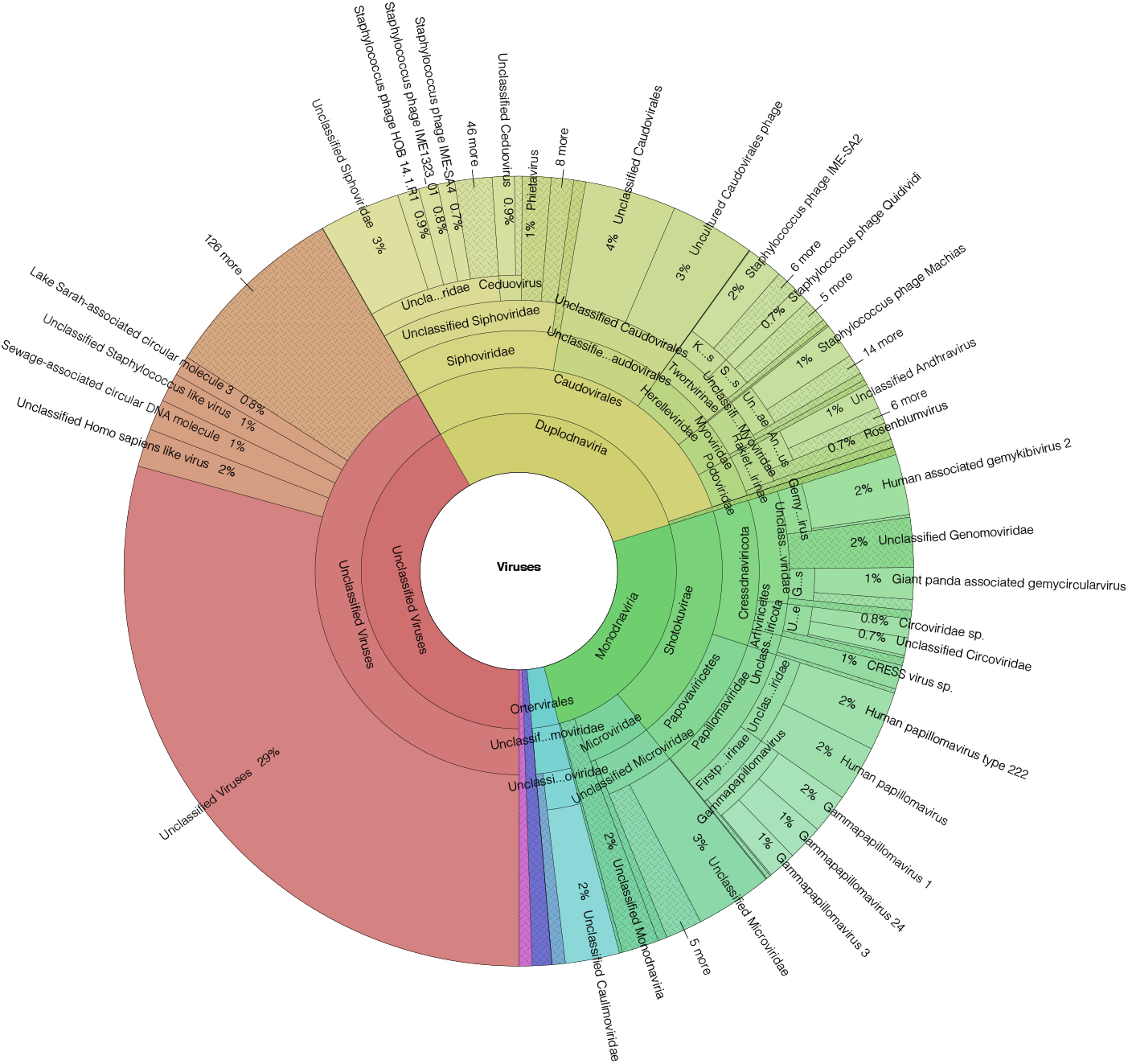
Overall distribution of viral taxonomy identified from 652 human skin virome metagenome samples derived from 42 subjects. Distribution consists of the total abundance of each taxonomic classification assigned to the assembled contigs >1000bp from human all skin metagenome samples that were identified as containing at least one viral gene using CheckV v.0.7.0 [39]. Double and single stranded DNA viruses composed more than 50% of the overall diversity abundance of the identified skin virome, however a large portion of highly abundant taxa were unable to be classified.

### 3.3 Subject Viral Diversity Abundance Across Skin Sites

The top ten most abundant viral families identified for each location across the entire group of participants are shown in Fig. 2. Of the most abundant viral families observed, *Papillomaviridae* and varying viral families that belong to the order *Caudovirales* were most abundant. This is consistent with the overall virome diversity identified across all individuals (Fig. 1). As seen in Fig. 2, the virome was different from subject to subject showing the individuality of the virome that could be used for human identification. When comparing across skin site locations within an individual, in some instances there was either a complete absence or addition of highly abundant viral families. One such family was the *Streptococcus satellite phage Javan 305*, although most of these other viruses have yet to be cultured or characterized. As for a comparison of the relative abundance of viral families, we observed clear differences across individuals. Considering the most abundant viral families, one could draw clear distinctions between individuals, while still maintaining higher levels of similarity within an individual across multiple skin collection locations. This observation further supports the notion that the human skin virome is an individual characteristic and potentially useful for discrimination between differing subjects.

**Fig. 2.**
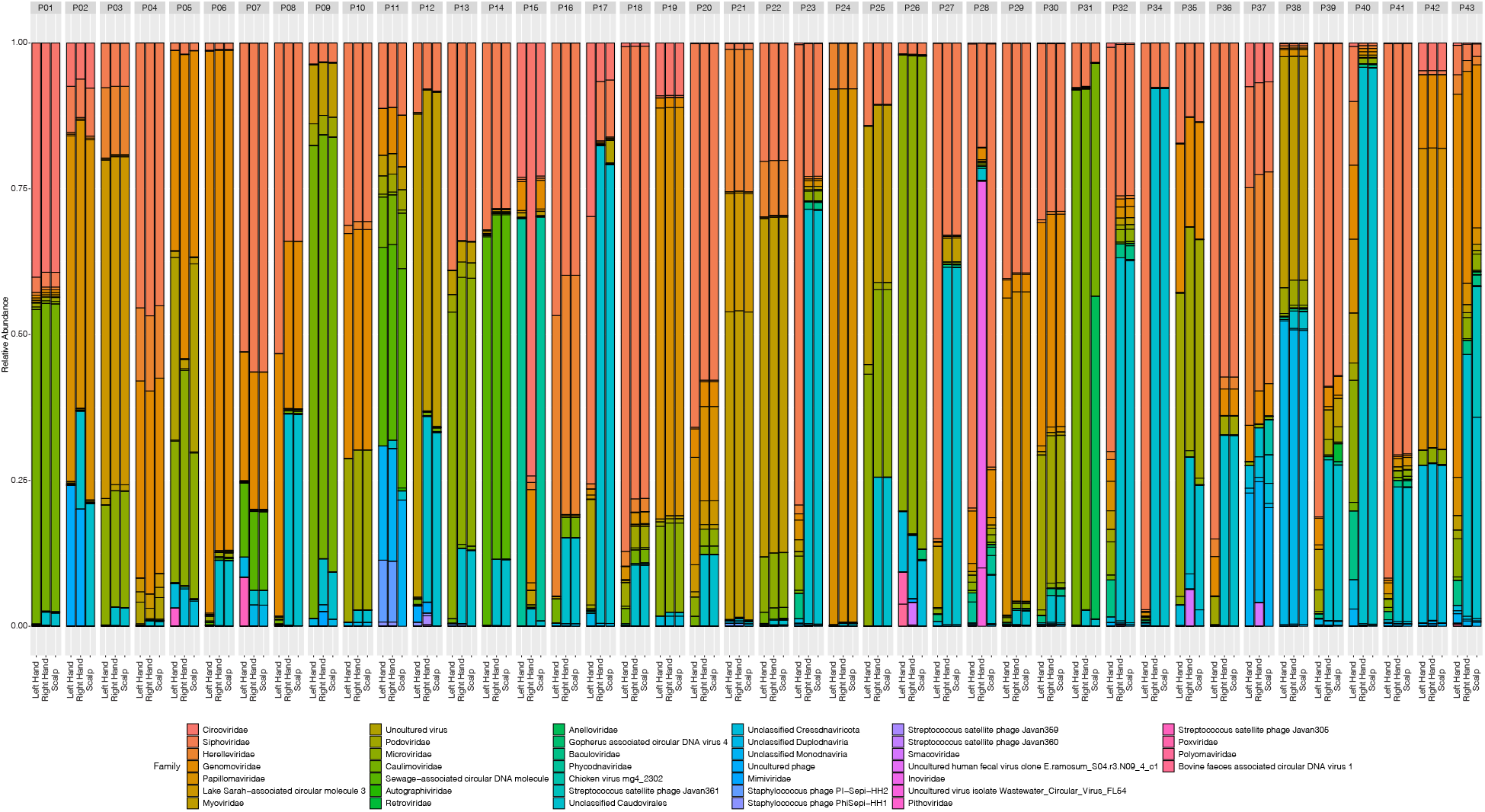
Relative abundance of the top ten most abundant identified viral families per subject by location shows inter-subject similarity, with increased between-subject differences in diversity. Of the identified viral contigs, abundance of all taxonomically assigned viral families were summed across all time points by anatomical sampling location within a subject. Each bar represents a location within a subject, as indicated at the bottom. The bars are then further separated by subject as indicated by subject notation at the top. All contigs that were not able to be taxonomically classified at the family level using currency databases were not included. Clear similarities in taxonomic abundance across locations within an individual can be observed. However, when comparing across subjects, there is increased dissimilarity in relative abundance taxonomic profiles than that of within subject comparison across locations.

### 3.4 Stability of the Skin Virome Overtime

The most abundant viral families of the human skin virome displayed a higher level of variation between individuals than variation within a single individual across sample locations (e.g., hands and scalp). To use viral markers for human identification and as forensic evidentiary material, these markers need to be identified with accuracy and consistency over time allowing for comparison of viral profiles collected at different time points. Therefore, the prevalence of contigs we identified were evaluated across individuals and locations within an individual over time. Stable viral taxa were categorized by the presence of a viral contig or taxa in at least four out of the five time points for a given individual at a certain skin location. This analysis identified the viral families, *Baculoviridae*, *Genomoviridae*, *Herelleviridae*, *Myoviridae*, *Papillomaviridae*, *Podoviridae*, *Siphoviridae*, *Unclassified Caudovirales*, and *Unclassified Homo sapiens like virus* to be stable in at least one individual across all three skin locations and thus was considered as potential target viral families for further investigation to develop biological markers for human identification (Fig. 3). Although these viral families may have stability in certain individuals, they may be absent or temporal in other individuals, therefore exhibiting variation in virome stability across subjects.

**Fig. 3.**
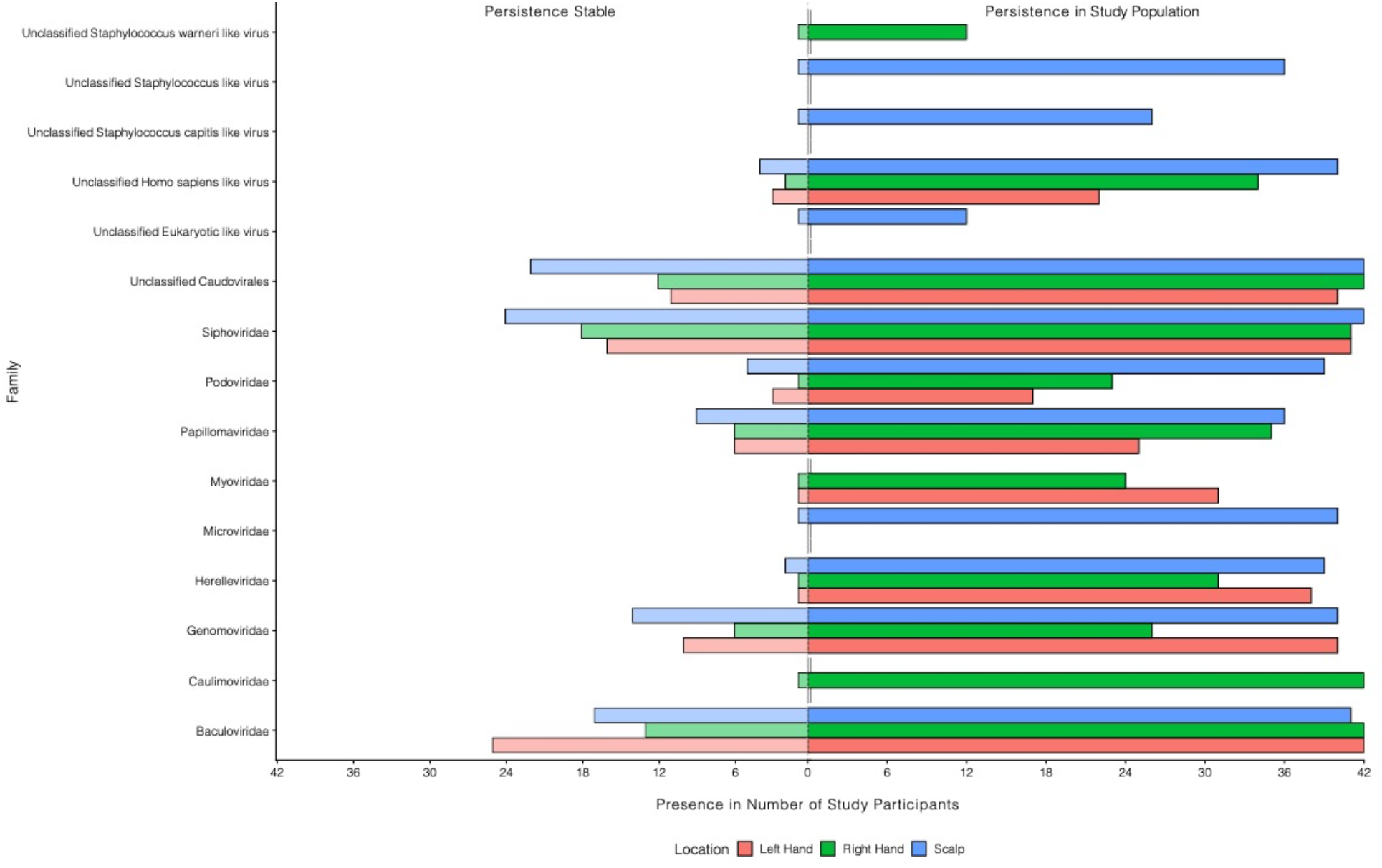
Persistence of stable identified viral families within the study population. Bars represent the number of study participants (42 total) within the study population where a denoted viral family was stable (left side; persistence stable) and the overall presence regardless of stability (right side; persistence in study population). Prevalence for each identified viral family was determined across the five sampled time points of initial sampling (day 0) and 2-week, 1-month, 3-months, and 6-months post initial sampling. Viral families that were present in four out of the five time points at an anatomical location within a subject were considered stable. Only viral families that exhibited stability in at least one individual within the population are displayed.

### 3.5 Identification of Putative Viral Skin Markers for Human Identification

Viral families alone cannot be utilized as a taxonomic marker for human identification and a higher level of annotation must be used (e.g., such as that of genus or species or unique contigs). Therefore, of the classified contigs, marker identification and stability were not only assessed at a family level but also at the level of genus and species resulting in identification of eight left hand, 11 right hand, and 23 scalp associated stable viral species (Table 1, SetA; Fig. 4A).

**Fig. 4.**
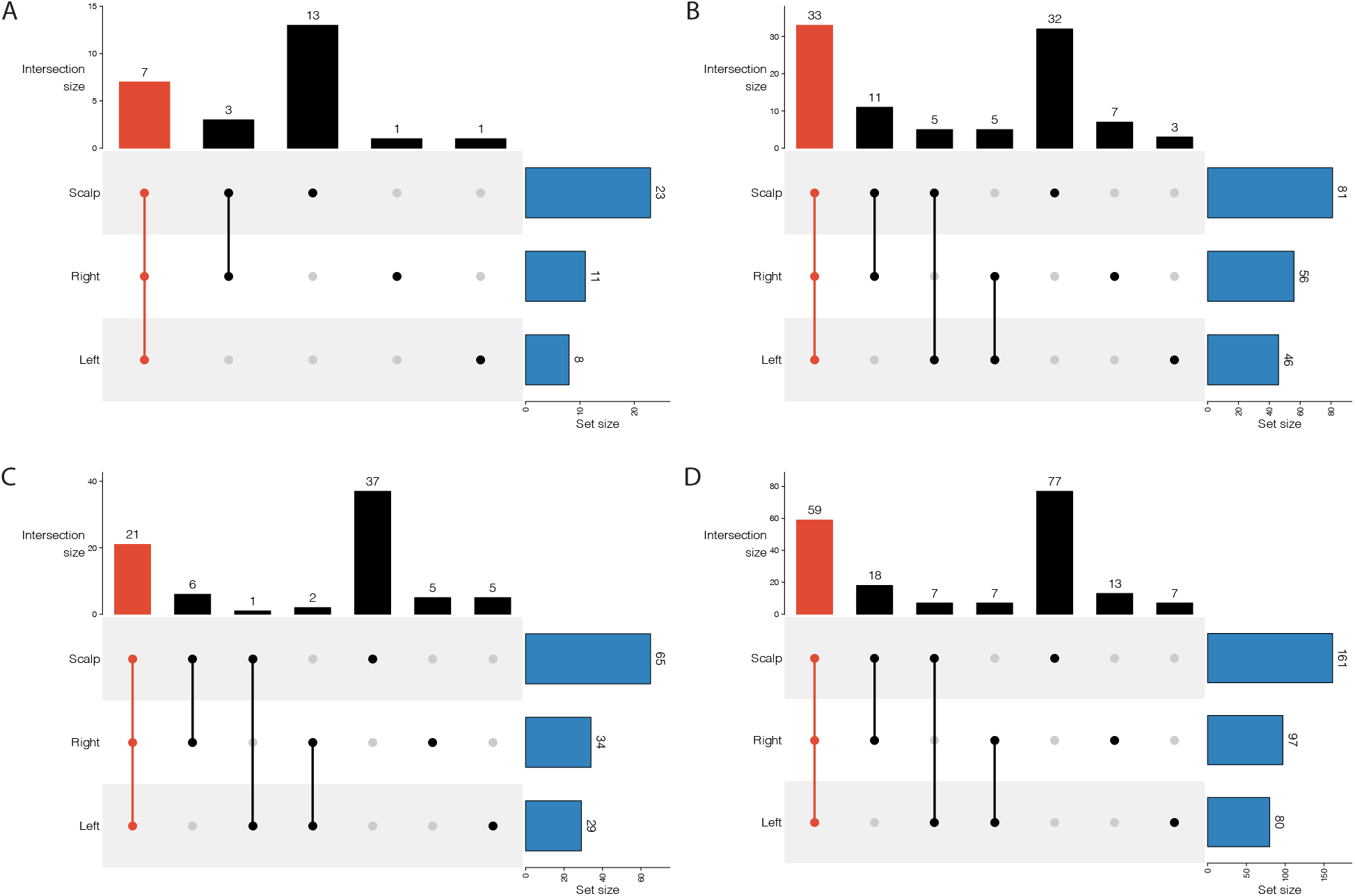
Prevalence of marker stability within the marker sets across the three anatomical locations. Plots show the distribution of viral markers that are stable in the study population across the three autosomal locations for Set A **(A)**, Set B **(B)**, Set C **(C)**, and all nonoverlapping markers from the three sets combined together **(D)**. Stability was defined as the marker being present in four out of the five time points within at least one subject at the denoted location. Highlighted in red is the quantity of markers that were deemed to be stable in all three autosomal locations and thus used for downstream statistical evaluation of population marker profile comparisons. Bars on top (Intersection Size) represent the number of markers that were considered to be stable at the combination of locations as noted by the scatter plot below the bar. Bars shown in blue (Set size) are the overall number of markers within the set that were found to be stable at each autosomal location. The 59 markers that were deemed to be stable across all three locations from all three sets combined **(D)**, are considered to be the putative human virome profile markers for human identification. Set composition is described in Table 1.

**Table 1.**
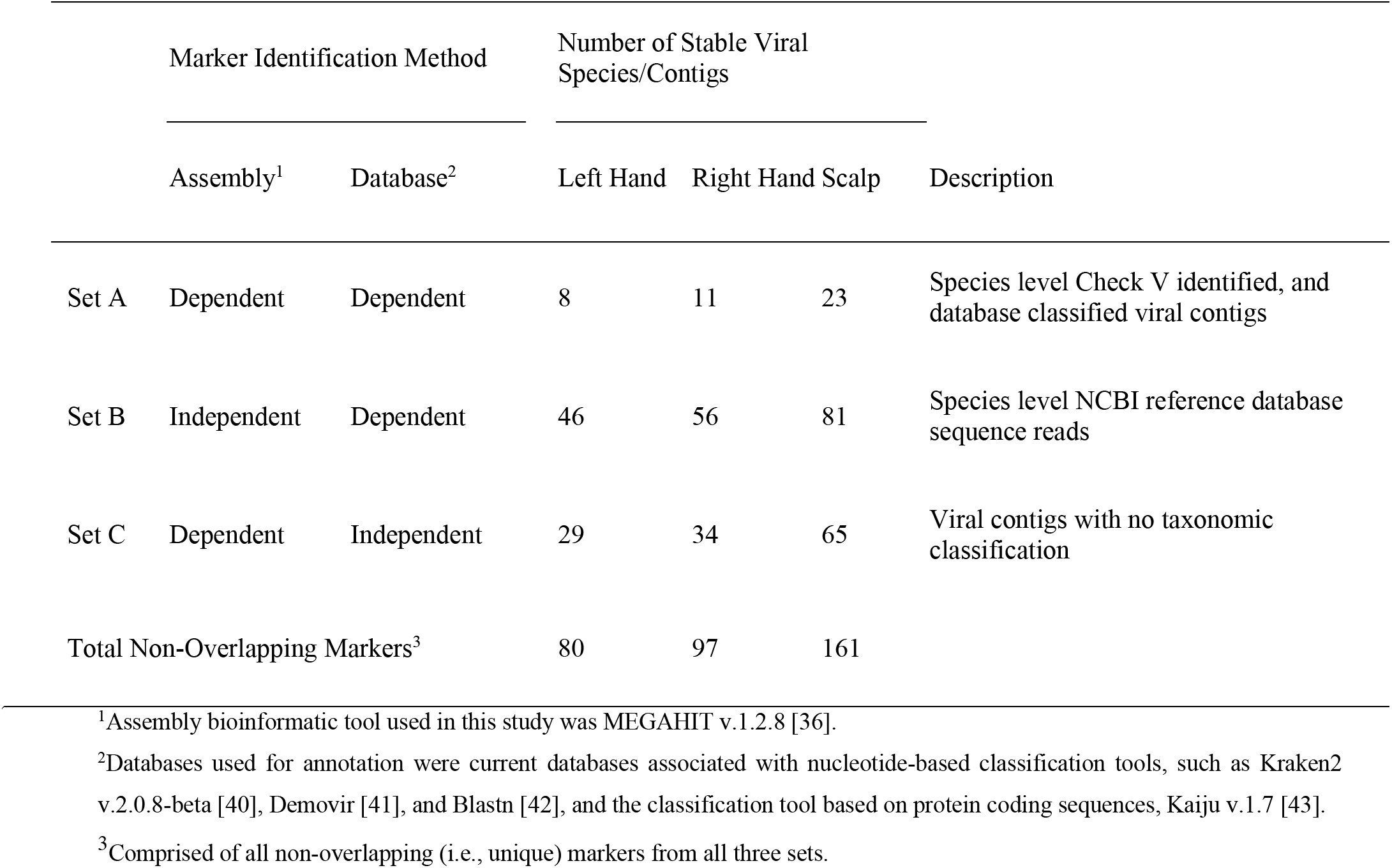
Description and quantity of the three human skin viral biomarker sets for human identification.

To improve viral genome recovery and reduce metagenome assembly bias, trimmed and quality filtered reads were mapped to all NCBI nucleotide viral reference sequences associated with target viral families identified to be stable across all three skin locations. All NCBI nucleotide viral sequences associated with the order of *Caudovirales* were used for this analysis due to the fact that several of the target families, such as *Siphoviridae*, *Podoviridae*, and *Myoviridae*, all fall under this order. Mapped counts to the reference genomes were evaluated similarly to that of classified viral contigs with species level analysis as opposed to family level evaluation performed previously. In addition, to identify candidate species or groups of species for human identification, virome stability within an individual over time was used as an important factor. Viral species found to be stable in four out of the five time points within a location for at least one individual were considered to be a stable viral species. In total, 46 left hand, 56 right hand, and 81 scalp associated stable viral species, using assembly independent and database dependent mapping, were identified as putative viral markers (Table 1, SetB; Fig. 4B).

To address viral dark matter that was unable to be annotated and viral identification bias based on only using known viral genes, stability of all contigs that were not identified as containing at least one viral gene using CheckV were assessed for stability within a subject. The subsequent contigs identified within a subject as being stable were retained and NCBI’s Blastn algorithm was used to annotate these contig sequences. Contigs containing <70% identity to a known organism or contigs with open reading frames with no synteny to prokaryotic or eukaryotic genes were considered to be putative viral sequences although viral origin could not be fully confirmed. For database independent marker identification, 29 left hand, 34 right hand, and 65 scalp contigs were identified as being stable (Table 1, SetC; Fig. 4C).

To address database bias, viral dark matter, and metagenomic assembly error, the three sets of putative viral markers for human contamination were compiled. When all three sets are taken into account a total of 188 non-redundant viral species or contigs markers were identified as stable and are thus proposed as potential viral markers for human identification (Fig. 4D). To further limit the number of markers, only markers that were found to be stable across all three locations within at least one individual were retained for marker identification, resulting in a final set of 59 putative human identification viral markers (Fig. 4D). A heatmap was rendered to visualize marker prevalence profiles of the 59 viral markers across the five time points by subject by location as shown in Fig. 5. Of the viral markers, seven markers (*Staphylococcus phage vB_SauH_DELF3*, *Unclassified Baculoviridae*, *Escherichia virus Lambda*, *Autographa californica multiple nucleopolyhendrovirus 1*, *Streptococcus phage phi-SC181*, *Marine virus AFVG_25M557*, and *Streptococcus phage phiJH1301-2*) were stable and present across all individuals, whereas all other markers retained discriminatory power across individuals.

**Fig. 5.**
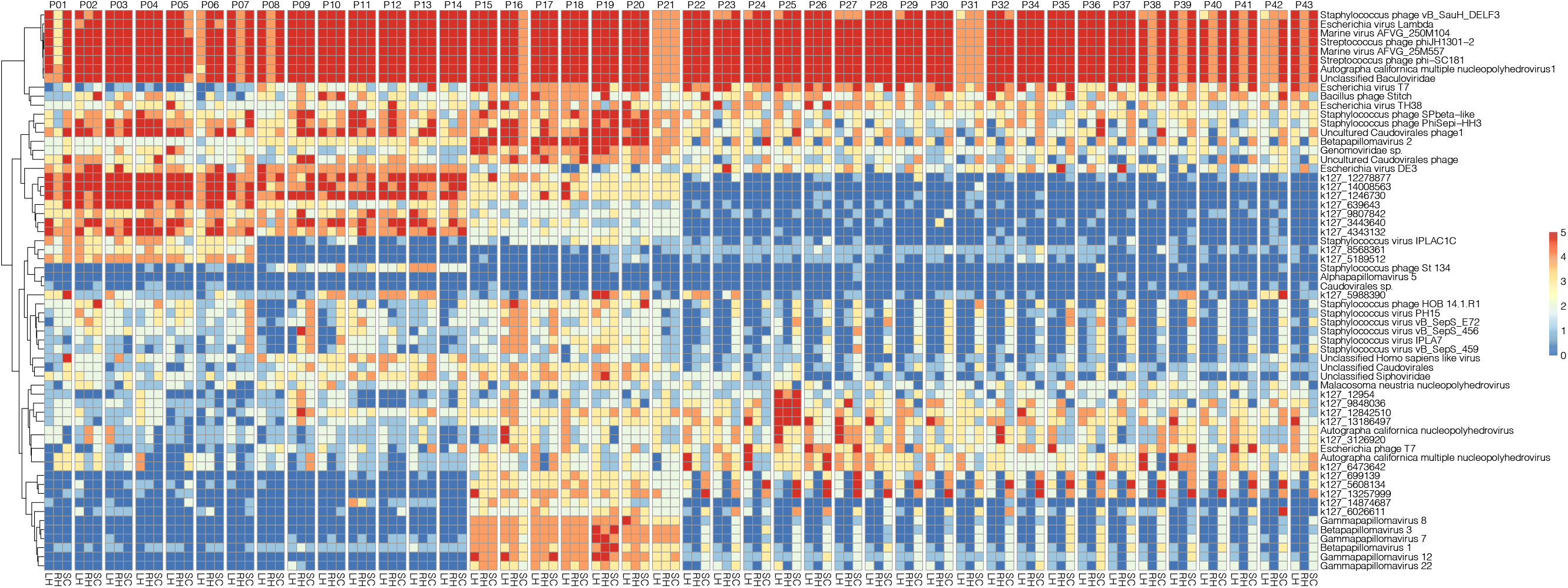
Prevalence of the 59 markers across the subjects and locations shows distinct profiles for each individual. Profile heatmap where each column is an anatomical location (LH = Left Hand; RH = Right Hand; SC = Scalp) which is separated by subject as denoted on top. Each row is associated with an identified viral stable marker from the overall marker set with all three sets (e.g., Set A, Set B, and Set C) combined. Rows are clustered based on marker prevalence similarity across the samples. Prevalence of the marker in the five time points is represented by the color scale with red being present in five out of the five time points and blue being present in zero out of the five time points. The top seven markers in the heatmap were found to be stable (stability defined as prevalent in four out the five time points) across all subjects and thus proposed for future studies looking at genetic variation within that viral species for subject discrimination within the population.

### 3.6 Statistical Assessment of Identified Viral Skin Markers for Human Identification

Three sets of putative markers for human identification were identified as described previously. These markers were chosen on their basis for stability across all sampled skin site locations within at least one individual thus proving their stability within one individual in the study population. However, in order to test the viability of the identified markers, the stability of the markers across the entire population and their differentiation across individuals must be statistically evaluated. To do so, for each marker set (which we labeled as “Set A”, “Set B”, “Set C”, and “Overall” which included all three sets combined) a binary Jaccard dissimilarity matrix was produced to compare within subject versus between subject variation on the basis of presence or absence of each of the markers within each set. For all three sets, within variation was significantly less dissimilar than that of the between variation (Set A: *P*=0.00011; Set B: *P*=4.4×10^-10^; Set C: *P*=1.5×10^-10^) (Fig. 6A-C). In order to evaluate the sets in combination with one another, we compared within subject variation and between subject variation using all markers for each skin site location. We additionally compared an overall set which encompassed all locations and subjects. For all site locations and the overall comparison (all locations and overall marker set comparison), there was a high significant difference between within subject variation and between subject variation (Left hand: *P*=6.3×10^-14^; Right hand: *P*<2.22×10^-16^; Scalp: *P*=3.6×10^-10^; Overall: *P*=5.3×10^-15^) (Fig. 6D-G). Thus, showing that the marker sets, on the basis of presence and or absence, are significantly more similar within an individual across time points than compared to the base-line presence and or absence of those same markers in the rest of the subject population.

**Fig. 6.**
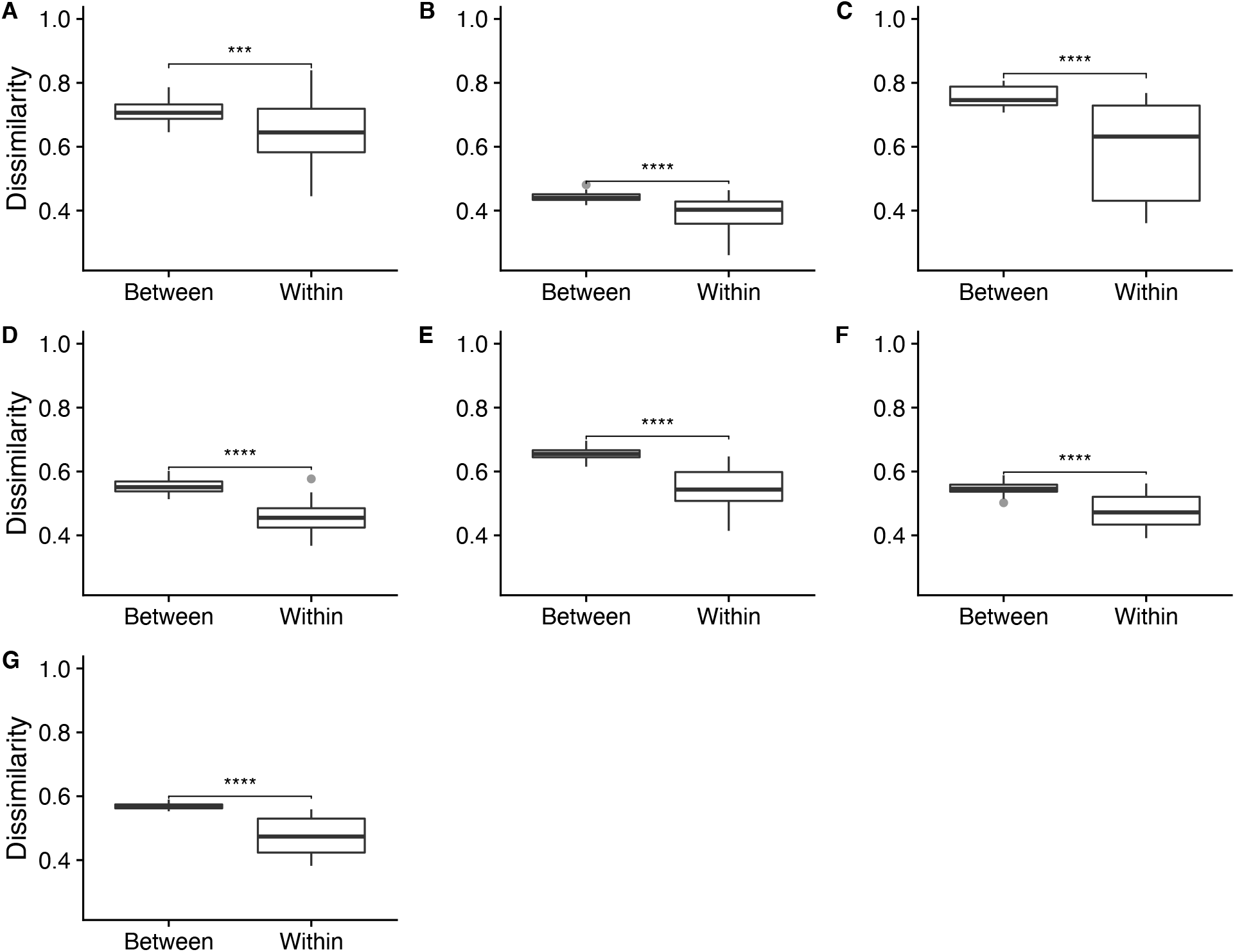
Profiles of identified viral biomarkers are stable over time as indicated by a significant decrease in inter-subject dissimilarity as compared to between subject dissimilarity. Boxplots of within-subject (comparison of samples within subject across time) and between (comparison of sample to all other subject samples) subject binary Jaccard dissimilarity distances; 0 = samples are identical, 1 = samples are disjoint. Statistical comparison of between-subject dissimilarity and within-subject dissimilarity was done by Wilcoxon ranked sum tests. Between-subject marker presence and absence profile was significantly more different than within-subject across time for the identified marker Set A (***; P<0.00011; *n* = 42 subjects) **(A)**, Set B (****; P<4.4×10^-10^; *n* = 42 subjects) **(B)**, and Set C (****; P<1.5×10^-10^; *n* = 42 subjects) **(C)**. When all non-overlapping identified stable markers from the three sets combined, between-subject marker presence and absence profile was significantly more different than within-subject across time for the anatomical locations left hand (****; P<3.6×10^-10^; *n* = 42 subjects) **(D)**, right hand (****; P<2.2×10^-16^; *n* = 42 subjects) **(E)**, scalp (****; P<6.3×10^-14^; *n* = 42 subjects) **(F)**, and overall all locations combined (****; P<5.3×10^-15^; *n* = 42 subjects)**(G)**. Description of composition of marker Set A, Set B, and Set C are described in Table 1. * significant at *P*<0.05; ** significant at *P*<0.01; *** significant at *P*<0.001; **** significant at *P*<0.0001

In order to evaluate subject differentiation across the population, the differences in marker diversity, using presence and absence of the identified markers, was evaluated. We compared within sample diversity (ɑ-diversity) of all of the markers using the Shannon index as a diversity metric (Fig. 7A). Using an ANOVA test, we found that there was a significant difference in the amount of marker diversity across subjects (**, *P*=0.002) and a slight significance difference across subject-reported gender (*, *P*=0.02). However, there was not a significant difference in ɑ-diversity across the three locations within subject (*P*=0.066). To establish if between subject-to-subject diversity (β-diversity) was significant, the Adonis test in the R package Vegan v.2.5-7 [50] was used to run PERMANOVA tests using Jaccard distance method with binary = True in order to evaluate differences on the basis of presence and absence and not abundance (Fig. 7B). It was found that for β-diversity there was a significant difference across subjects (*P*<0.001) with an effect size of R^2^ = 0.3415.

**Fig. 7.**
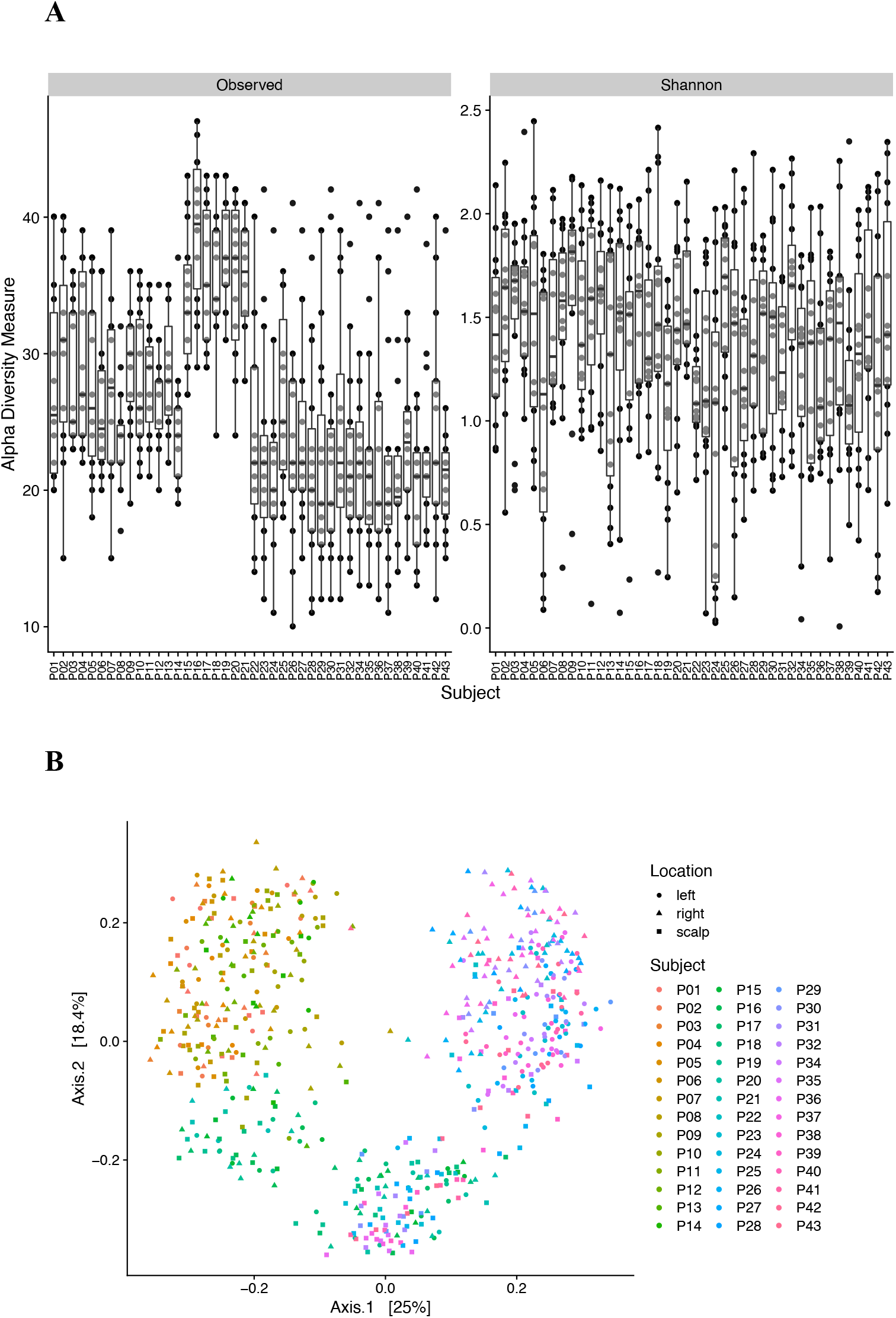
Identified viral marker diversity across subjects. **(A)** Boxplots comparing ɑ-diversity of the identified viral markers in subjects using the ɑ-diversity measure Shannon [49], as well as the uncorrected number of observed markers within subject. An ANOVA for the variables of subject (**, *P*=0.002) and gender (*, *P*=0.02) were found to be significant, while location within subject was not (*P*=0.066). **(B)** PCoA plot of binary Jaccard dissimilarity distances of virome samples by subject to assess β-diversity of the identified viral markers across subjects. Clustering by subject was significant (***, *P<*0.001) by PERMANOVA analysis using the Adonis test with 34.15% of the variation explained by subject (R^2^ = 0.3415).

## 4. Discussion

Utilization of alternative sources of biological samples for human identification must incorporate aspects of both target marker stability over time and the ability to utilize those markers to have probative discriminatory power across individuals within a population. This study set out to address the viability of the human skin viral microbiome (virome) for its stability and utilization as a source for human identification. Previous studies have shown certain viral taxa associated with human skin disease to have yearlong stability [54]. However, no previous published studies have investigated the potential utilization of the skin virome for forensics or to identify potential target viral markers for human identification.

In this study, we sequenced a total of 652 human skin viral metagenomes and analyzed that data to identify viral markers for human identification. The sequencing data was assembled into viral contigs. Of the 62,101 assembled contigs with >1000 bp, 1298 contigs were identified as viral contigs based on current database dependent identification tools. This is a small percentage of the overall metagenome assembly. To ensure that the viral contigs were devoid of bacterial contamination, the contigs were analyzed for the presence of 16S rRNA genes. The assemblies contained on average 0.002% of ribosomal reads per sample with a maximum of 0.16% of ribosomal reads. Previous studies have demonstrated that if viral metagenomes have less than 0.2% 16S rRNA reads that these datasets are enriched for viral sequences and have minimal and likely negligible bacterial contamination [52]. Therefore, the low occurrence of bacterial 16S rRNA sequences suggests our viromes to be highly enriched for human-associated viruses with minimal contamination. Since viral genes within contigs were determined by comparing to current databases, our data suggests that the current databases are lacking in viral diversity and this lack of representation of viral diversity in databases makes it difficult to study viromes in any environment. Thus, similar studies coupled with genome annotations are greatly needed to improve viral annotation. As such, this study should be viewed as a first step towards uncovering viral diversity and stability in the human virome in addition to its application to forensics.

Of the identified viral contigs, both double and single stranded DNA viruses were identified. The most abundant ds DNA viruses identified were that of bacteriophage belonging to the viral taxonomic order *Caudovirales*. Previous studies have found the skin virome diversity to be mainly composed of *Caudovirales* bacteriophage [28, 54] and our findings here were in agreement with this observation. Viral families identified in skin virome that fall under the order *Caudovirales* included *Siphoviridae*, *Podoviridae*, and *Herelleviridae*. These viral families include bacteriophage that are obligately associated with bacteria commonly found in the skin microbiome (i.e., *Staphylococcus* and *Streptococcus*) [28,55,56]. With bacteria predominating in the skin microbiome, it is logical to speculate the increased abundance of bacteriophages would follow. These bacteriophages may help control bacterial populations in the skin and may help structure bacterial communities in the human skin microbiome with relation to environmental and lifestyle of the person. In addition to double stranded DNA viruses, a large abundance of single stranded DNA viruses was also identified in the skin virome samples. Small circular DNA viruses, those that are typically associated with Eukaryotes, such as *Adenoviridae*, *Anelloviridae*, *Circoviridae*, *Herpesviridae*, *Papillomaviridae*, and *Polyomaviridae* have all previously been reported to be associated with the human skin virome. However, the identification of small Cress-like DNA phages has not previously been reported from the human skin. This is due to the recent discovery of novel Cress-DNA viruses and their recent addition to NCBI databases [28,53,55,57]. A large proportion of the single stranded viruses identified are Papillomaviruses, which are common skin associated viruses that can act as opportunistic pathogens [58].

We also observed a high abundance of the small circular single-stranded DNA *Cressdnaviricota* viruses across all subjects in this study. These viruses have previously not been reported at high abundance in the human skin virome. This finding could potentially be attributed to the recent identification of this novel small single stranded DNA virus group [57]. Of the contigs that were annotated under *Cressdnaviricota* many of them only displayed similarity to *Cressdnaviricota* viruses. However, these viruses were not identical to the sequences in the current database suggesting the viruses identified in this study could be novel viruses related to *Cressdnaviricota* and other small DNA viruses.

Many contigs that were identified as being of viral origin were small and circular in nature. These viruses have similarity to other small circular DNA viruses in current viral reference databases. However, due to their low percent identity to any known virus or organism they remained unclassified in our dataset at lower taxonomic levels (e.g., at the level of viral genus or species). This was especially evident in unclassified viruses having similarity to viruses belonging to family *Microviridae*. As previously mentioned, the skin virome contains many novel viruses, particularly noted here in the family *Microviridae*. In order to better characterize the viral diversity in the human skin virome, more research into viral discovery is needed. However, with a large proportion of viral contigs having no similarity to characterized viruses, we implemented a database independent approach paired with database dependent methods to alleviate annotation inaccuracies and missing reference material.

In order to evaluate the applicability of human virome for subject identification, viral contig diversity and abundance across locations and subjects was further analyzed to identify the top ten most abundant viral families for each subject by sampling location (Fig. 2). Distinct differences in the relative abundance of these contigs were observed across individuals. Some individuals had similar profiles (for example, the viral profiles of subject 21 (P21) and subject 38 (P38) were strikingly similar for unrelated and/or non-cohabitating subjects), however even within individuals of similar contig prevalence, the contig abundance was variable allowing for subject differentiation. In addition, when comparing subject to subject variation, relative abundance across locations within an individual was also visualized (Fig 2.). Similarity across all three locations was observed within most individuals. Differences were noted for some subjects between locations (right hand, left hand, and scalp). It is hypothesized that this may have to do with hand dominance coming into contact with that individual’s hair and scalp thus sharing similar viral taxa across locations. The main difference observed across these locations is either the addition of a viral family or the loss of one.

It is important to note, however, that by comparing just subject and location within subject, with time of sampling and viral stability not taken into account, we were able to identify the top ten most abundant taxa for each sampling anatomical location and subject summed across the five collection time points. In other words, the relative abundance of the most abundant annotated contigs displayed greater variation between subjects compared to that of within subject variation across sampling locations (Fig. 2). This suggests viral marker presence can be discriminatory for differentiation of individuals within the study population.

Not only do viral markers need to be diagnostic for individuals, but they also need to be stable over time. For forensic purposes, we identify that the core stable virome needs to be targeted for biomarker development. The virome, especially concerning viruses found on body locations that are constantly exposed and that interact with other objects and the environment, will potentially have a large temporal component in addition to the stable virome [28, 51]. Temporal viruses do have the potential to provide informative forensic information that would be unique to certain lifestyle characteristics like occupation, contact with animals, recent travel, or hobbies (such as gardening, etc.). Temporal viruses, while not stable for direct forensic applications, could provide important information towards identifying circumstantial characteristics specific to an individual. However, for the purpose of acting as an alternative genetic source to traditional STR methods, target viral markers must be stable over time, and therefore, we required that viral markers be present during our entire sample collecting regime, which we repeated for each individual at all three locations over a time course of 6-months.

For the purposes of this study, we considered a virus to be stable if a particular consistently annotated virus was present in at least four out of the five time points collected within a single anatomical location for an individual. Of the viral families identified in the overall assembly, 15 families were considered to have stability in at least one location within an individual. Of the 15 families, nine viral families presented stability in at least one individual for all three body locations sites. Of note, many of the nine families fell under the order of *Caudovirales*. As previously mentioned, many studies have evaluated and classified the core bacterial microbiome on the skin and thus it is not surprising that bacteriophage would be both present and temporally stable seeing as how their bacterial hosts are also present and stable on human skin [51,55,59]. In addition to families of *Caudovirales*, *Papillomaviridae* was also found to be stable which is consistent with findings observed in another study [54]. However, in a few older studies, Papillomaviruses were found to not be stable or not found to be as stable as the communities of certain phage viruses as well as both bacterial and fungal microbiome communities [28, 51]. Interestingly both *Baculoviridae* and *Genomoviridae* were also found to be stable across multiple individuals in all three physical locations we sampled. Notably, *Baculoviridae* are sometimes used for bioengineering purposes in sequencing laboratories so there is a chance that their presence in our study is due to contamination from a sequencing provider, however, we think this is highly unlikely as certain *Baculoviridae* have the ability to infect mammalian cells, such as Autographa californica multi-nuclear polyhedrosis virus. Therefore, our identification of similar viruses may potentially be a true biological observation that has not previously been accounted for due to other studies assuming it is contamination [60]. Although *Genomoviridae* have previously been identified from human vaginal samples, very few studies have explored these small single stranded DNA viruses and their diversity and stable presence as a part of the human skin core virome [61]. However, previous studies have identified *Genomoviridae* as being a stable and persistent virus in other mammals such as that of bats [62].

Of the nine viral families that we identified as being temporally stable, their continuously observed persistence was found across many, though not all, of the subjects in our study. This is potentially due to certain viral genera or species within a particular family as being temporally stable as opposed to similar genera or species within that same family that may not have been temporally stable. Additionally, there is a possibility that our viral metagenome assemblies could be biased due to the high amount of repetitive sequences seen in some particular viruses. Therefore, in order to reduce assembly-based bias and increase annotation, we trimmed sample reads and mapped that data to all genome sequences available in NCBI’s nucleotide database pertaining to the order *Caudovirales* and the viral families *Papillomaviridae*, *Genomoviridae*, and *Baculoviridae,* to improve viral detection and sample annotation. Species level mapped NCBI reference genomes were evaluated during this process. In addition to stability of reference mapped counts, species level stability was also assessed for viral contigs. Species that were identified as being persistent across all three locations in at least four out of the five time points for an individual were considered for putative viral markers for human identification.

In order to address viral diversity missing from databases, we first filtered sequencing reads on the basis of base-call quality and then mapped the sequencing reads to the total contigs in our viral metagenome assembly. Contig sequences that were stable within an individual for each sampling location were analyzed with Blast using the Blastn alignment algorithm. Contigs that did not share a high percent similarity (>70%) to known non-viral genomes were considered to be of potential viral origin and retained in the analysis. Of the contigs that were deemed to be of potential viral origin, those that were stable across all three body locations were considered to be additional putative viral markers for human identification, though their exact taxonomic classification is not known because they did not show synteny to any reference sequences.

In this study we identified a total 188 viral markers, which included all stable NCBI reference mapped species, stable viral contig annotated species, and stable potential viral contigs that showed limited synteny to databases or included viral-like gene regions. Of those 188 viral markers, 59 markers were found to be stable across all three body site locations sampled (Fig. 4D). These 59 target viral markers are proposed as having potential for the purpose of human identification, as shown across 42 individual test subjects. Of the 59 viral markers, seven were persistent across >90% of the sample population. These seven markers may not be usable for subject-to-subject individualization, in regards to their presence and absence in the virome of an individual due to their low power of discrimination. However, they are prime candidates for single nucleotide variation comparison or RFLP analysis across a sampled population which we suspect will add to an additional level of marker exploitation at the scale of single nucleotide variation, as well as presence/absence of insertions and deletions at the nucleotide level. As for the other 52 markers, they may be used as a presence/absence basis for human identification.

Distinct profile patterns were observed across the subjects (Fig. 5) for specific viral taxa. In particular, for subject participants P15-P21, they share similar presence/absence profiles for certain viruses such as that of *Gammapapillomaviruses* which were not observed in the rest of the population. The presence of these district profile subpopulations is not due to sample processing or sequencing contamination due to the randomization of processing of samples and is believed to be a true biological finding. However, it is noted that for the subpopulation of P15-P21 these subjects, frequently shared physical space and objects or were a cohabitating partner of one of the members of the aforementioned group. Not only does this subpopulation stand out from the rest of the population, but they can also be distinguished from one another based on their profiles, thus adding a potential additional level of discrimination. Suggesting that not only can the virome be used as an individualizing characteristic but also bares circumstantial lifestyle or environmental characteristics. This data indicates that additional work is merited to examine subpopulations of people that cohabitate or share working environments.

The identified markers were statistically evaluated and were found to be more significantly similar within subjects across time points compared to between subjects using presence/absence of the three identified marker sets. The more markers that were used (i.e., the combination of all sets) resulted in a greater significance in the similarity differences (*P*=5.3×10^-15^). Thus, the addition of sets B and C were necessary in development of a more stable set for human identification profile production and evaluation. Diversity across subjects was also evaluated. We found that subject to subject variation was a significant variable associated with both ɑ-diversity and β-diversity. Therefore, showing that not only are these viral markers stable but there is significant difference in subject-to-subject diversity which highlights the ability to separate the skin virome of one individual from another.

## 5. Conclusions

Viral biomarkers were identified from human skin virome metagenome samples from 42 individuals with the goal of developing and assessing the potential for human identification and diagnostics. In total, we identified 59 putative markers and we found seven markers that were present across all subjects and have potential to be used as targets for future studies into SNP and genetic variation within target viruses that could be used for discrimination of individuals within the population. We found the remaining 52 markers, when taken as a total community on the basis of presence and absence, were statistically different for subjects and thus act as a full set of markers for human identification profile production.

## Data and Software Availability

All raw sequencing data has been deposited in the NCBI Short Read Archive (SRA) under the accession code PRJNA754140. All metadata, list of markers, contig sequences, annotation files, and scripts described in the material and methods are publicly available and archived at: https://github.com/HerrLab/Graham_2021_forensics_human_virome.

## Acknowledgments

We thank all participants for their contribution and participation in this study. We also thank W. Tom, A. Neujahr, N. Aluthge, C. Anderson, and others in the Fernando Lab who assisted with training of experimental methods and bioinformatic support. This work was completed using the Holland Computing Center of the University of Nebraska, which receives support from the Nebraska Research Initiative.

## Author Contributions

Author contributions: MA, JC, SF, and JH contributed to the experimental design and conceptualization of the study; MA was responsible for participant recruitment and sample and metadata collection; EG and SF developed virome processing methodology and prepared samples for sequencing; EG and JH developed the bioinformatic pipeline and analyzed sequence data; JC contributed statistical support; EG, MA, JC, SF, and JH drafted the manuscript.

---

**Supplementary Fig. 1.**
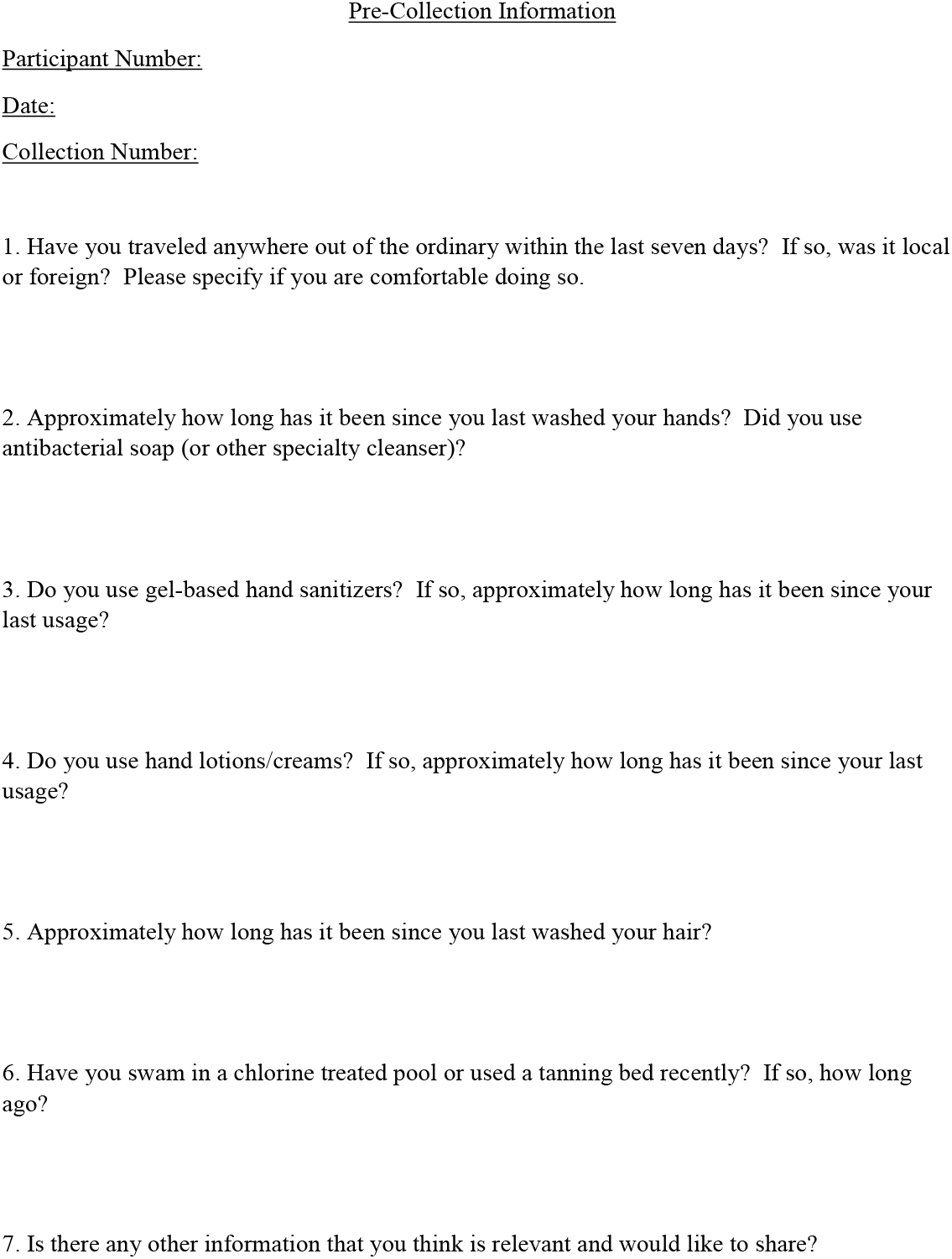
Pre-Collection questionnaire given to all participants prior to each sampling time to collect pertinent metadata. Questions were designed to address potential sources of variance that may affect viral diversity on the hands and the scalp. Participants were given a de-identified participant code (P#) at the initial date of sampling and that code is what was referred to for the entirety of sample processing, analysis, and what was reported and referred to in this study. All metadata collected from this questionnaire is publicly available at: https://github.com/HerrLab/Graham_2021_forensics_human_virome.

**Supplementary Fig. 2.**
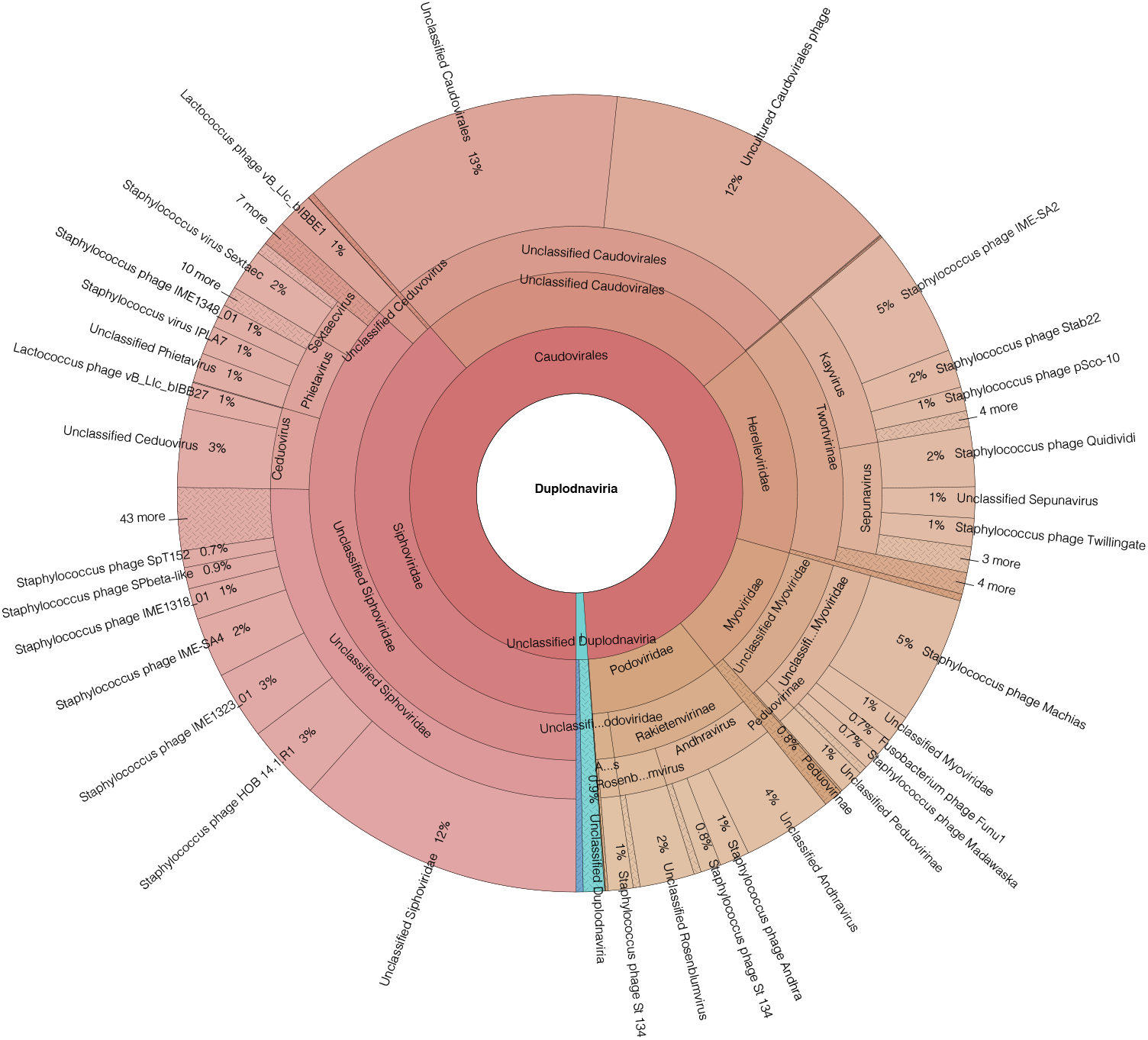
Distribution of the taxonomy under the viral realm *Duplodnaviria* (i.e., double stranded DNA viruses) identified from 652 human skin virome metagenome samples. Distribution consists of the total abundance of each taxonomic classification assigned that were classified in the realm of *Duplodnaviria*. Viruses that were annotated were classified from assembled contigs >1000bp from human all skin metagenome samples that were identified as containing at least one viral gene using CheckV v.0.7.0 [39]. *Duplodnaviria* composed 29% of the overall diversity abundance of the identified skin virome (Fig. 1).

**Supplementary Fig. 3.**
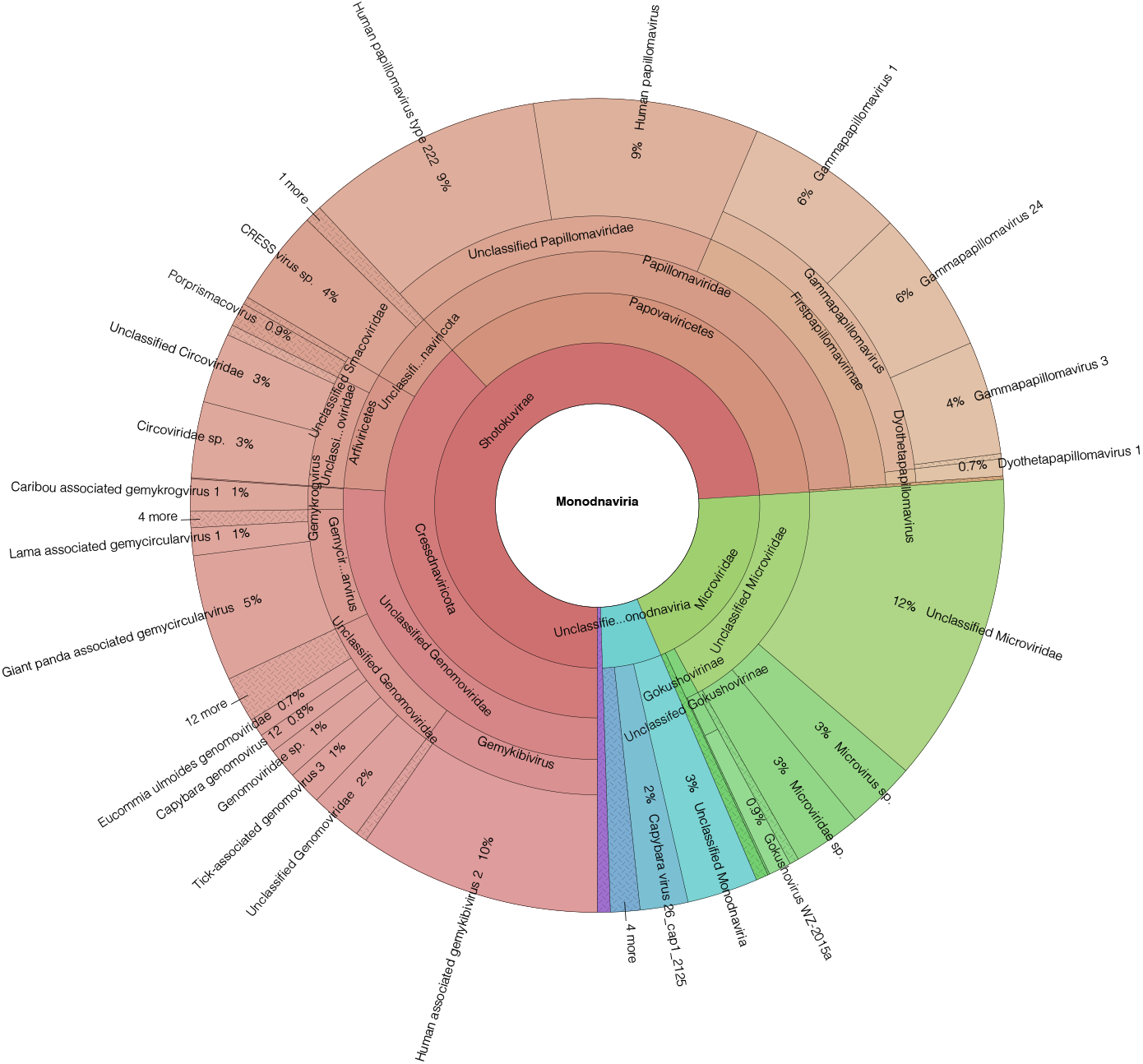
Distribution of the taxonomy under the viral realm *Monodnaviria* (i.e., single stranded DNA viruses) identified from 652 human skin virome metagenome samples. Distribution consists of the total abundance of each taxonomic classification assigned that were classified in the realm of *Monodnaviria*. Viruses that were annotated were classified from assembled contigs >1000bp from human all skin metagenome samples that were identified as containing at least one viral gene using CheckV v.0.7.0 [39]. *Monodnaviria* composed 26% of the overall diversity abundance of the identified skin virome (Fig. 1).

